# An individual’s skin stiffness predicts their tactile acuity

**DOI:** 10.1101/2023.07.17.548686

**Authors:** Bingxu Li, Gregory J. Gerling

## Abstract

Individual differences in tactile acuity have been correlated with age, gender, and finger size, while the role of the skin’s stiffness has been underexplored. Using an approach to image the 3- D deformation of the skin surface while in contact with transparent elastic objects, we evaluate a cohort of 40 young participants, who present a diverse range of finger size, skin stiffness, and fingerprint ridge breadth. The results indicate that skin stiffness generally correlates with finger size, although individuals with relatively softer skin can better discriminate compliant objects. Analysis of contact at the skin surface reveals that softer skin generates more prominent patterns of deformation, in particular greater rates of change in contact area, which correlate with higher rates of perceptual discrimination, regardless of finger size. Moreover, upon applying hyaluronic acid to soften individuals’ skin, we observe immediate, marked and systematic changes in skin deformation and consequent improvements in perceptual acuity. Together, the combination of 3- D imaging of the skin surface, biomechanics measurements, multivariate regression and clustering, and psychophysical experiments show that subtle distinctions in skin stiffness modulate the mechanical signaling of touch and shape individual differences in perceptual acuity.

**Key points described in the manuscript:**

- While declines in tactile acuity with aging are a function of multiple factors, for younger people the current working hypothesis has been that smaller fingers are better at informing perceptual discrimination due to a higher density of neural afferents.
- To decouple relative impacts on tactile acuity of skin properties of finger size, skin stiffness, and fingerprint ridge breadth, we combined 3D imaging of skin surface deformation, biomechanical measurements, multivariate regression and clustering, and psychophysics.
- The results indicate skin stiffness generally correlates with finger size, although more robustly correlates with and predicts an individual’s perceptual acuity.
- In particular, more elastic skin generates higher rates of deformation, which correlate with perceptual discrimination, shown most dramatically by softening each participant’s skin with hyaluronic acid.
- In refining the current working hypothesis, we show the skin’s stiffness strongly shapes the signaling of touch and modulates individual differences in perceptual acuity.

## INTRODUCTION

Tactile acuity varies among individuals and declines over one’s lifespan, as influenced by finger size, skin stiffness, age, gender, and cognitive factors. Declines related to aging intertwine several causes (Montagna & Carlisle, 1979; Gescheider *et al*., 1994; Stevens & Patterson, 1995; Stevens & Choo, 1996; Deflorio *et al*., 2023). In contrast, in younger age cohorts, finger size alone has been correlated with acuity (Peters *et al*., 2009; Peters & Goldreich, 2013; Abdouni *et al*., 2018; Olczak *et al*., 2018). The current working hypothesis is that the density of neural afferents determines perceptual acuity. Given that individuals generally have the same number of afferents, smaller fingers lead to higher afferent density and superior ability (Peters *et al*., 2009). Therefore, women, due to smaller fingers on average, tend to demonstrate higher tactile acuity than men. However, as noted in Peters, et al. (2009), the observed inter-individual variance in tactile acuity remains yet to be fully explained even upon accounting for finger size.

In addition to one’s finger size, an individual’s skin mechanics may influence his or her tactile acuity. For instance, in cases of age-related degradation, decreased tactile sensitivity has been observed with changes in skin mechanics, apart from changes in cognitive functioning (Wickremaratchi & Llewelyn, 2006; Bowden & McNulty, 2013; Pleger *et al*., 2016). Indeed, such reductions in tactile acuity may result from a loss of elastin fibers and increased wrinkling of the skin (Abdouni *et al*., n.d.). Moreover, aging women tend to experience decreased skin thickness, which also reduces tactile acuity, and in a much more dramatic way than for aging men (Abdouni *et al*., n.d.), who exhibit larger and thicker finger pads, due in part a thicker stratum corneum (Peters *et al*., 2009). Furthermore, and most evident when isolated in younger people in their twenties and thirties, increased skin conformance (i.e., a measure of how much the skin invades the gaps in grating stimuli) is associated with lower tactile detection thresholds (Woodward, 1993; Vega-Bermudez & Johnson, 2004; Gibson & Craig, 2006). Some have suggested that increased skin conformance may increase neural firing rates of afferents responding to edge stimuli (Phillips & Johnson, 1981), though the results remain inconclusive (Hudson *et al*., 2015). How such changes in the skin’s mechanics influence perceptual discrimination remains unanswered. In addition to skin stiffness, fingerprint ridge breadth may play a role. In particular, the dense papillary ridges of the skin are filled with sweat ducts and sites of afferent innervation (Loesch & Martin, 1984*a*, 1984*b*; Dillon *et al*., 2001; Abdouni *et al*., n.d.) while the underlying and opposing intermediate ridges are sites where changes in the stiffness of skin layers may amplify the response sensitivity of originating afferents, especially for slowly adapting type I fibers (Gerling & Thomas, 2008; Gerling, 2010; Pham *et al*., 2017).

Herein, we seek to evaluate the impact of the skin’s stiffness and fingerprint ridge breadth on individual differences in perceptual acuity in younger participants. This effort seeks to evaluate the current working hypothesis that, for younger people, those with smaller fingers demonstrate higher tactile acuity. Therefore, we perform a series of biomechanical, psychophysical, and statistical modeling experiments to decouple relative impacts on tactile acuity of skin properties, including finger size, skin stiffness, and fingerprint ridge breadth. In analyzing these factors, 3-D stereo imaging is used to capture the deformation of the skin surface over the time course of stimulus indentation, perceptual discrimination is evaluated across a robust and naturalistic range of object compliances, and hyaluronic acid is used to soften the skin to directly modulate its stiffness for perceptual evaluation.

## RESULTS

A series of five experiments are performed. First, relationships are evaluated between measurements of finger size, skin stiffness, and fingerprint ridge breadth, and each in turn with perceptual discrimination. Second, imaging is used to characterize patterns in the deformation of the skin’s surface, using a 3-D stereo technique. Third, measurements of skin deformation indicate how respective finger properties modulate levels of discrimination. Fourth, statistical models are used to decouple finger size and skin stiffness and characterize the performance of distinct subgroups from among the 40 participants, which do not divide entirely upon gender lines, but are driven more by an individual’s skin stiffness. Finally, hyaluronic acid is applied to soften the skin stiffness of a new cohort of participants to evaluate before and after perceptual performance.

### Experiment 1: Relationships between finger properties and perceptual discrimination

Perceptual discrimination of material compliance is evaluated in passive touch via pairwise comparison. Four compliance pairs are used, 45/10, 184/121, 33/5, and 75/45 kPa, Fig. 1(a). The 45/10 kPa pair is readily discriminable by all participants, while the 75/45 kPa pair is not discriminable at a threshold of 75% correct. Between these extremes, the 33/5 and 184/121 kPa stimulus pairs are discriminable. Note the compliance values of these pairs were selected because they span either side of the skin’s stiffness, observed to be 42 - 54 kPa (Miguel *et al*., 2015; Gerling *et al*., 2018).

**Figure 1.**
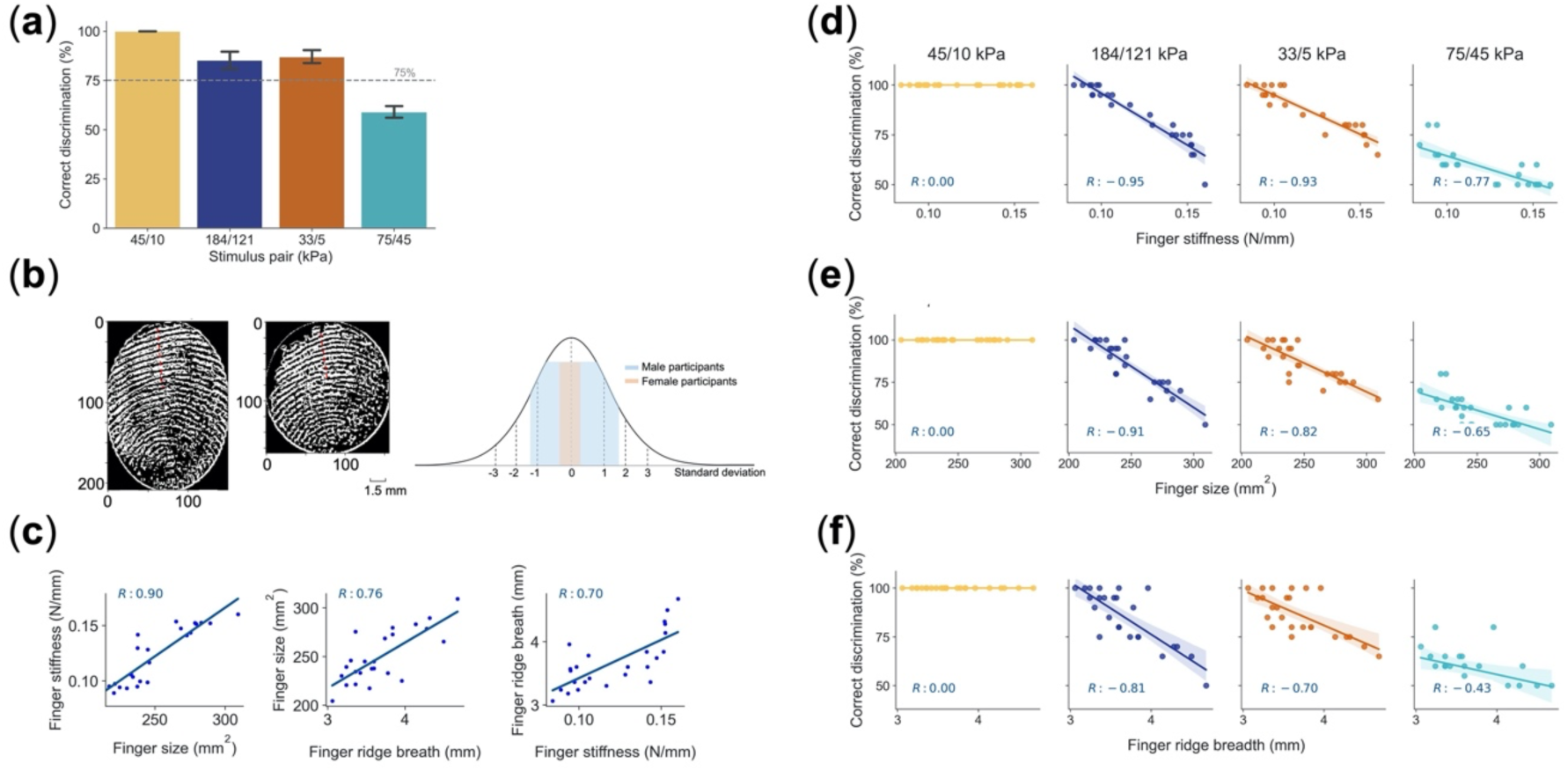
Relationships between finger skin properties and perceptual discrimination. (a) Pairwise discrimination of compliant stimuli indicates 45/10 kPa pair is readily discriminable, 184/128 and 33/5 kPa pairs are discriminable above 75% threshold, and 75/45 kPa pair is not discriminable. (b) Fingertip contact with glass plate for two participants to measure ridge breadth. Overlay of finger widths of this study’s participants (colored segments) over dataset of 4,000 people (Tilley & Associates, 2001). Our study’s male participants span more than one standard deviation, while female participants exhibit less variance. (c) Relationships per participant between finger size, stiffness, and ridge breadth, indicating highest correlation between finger stiffness and size. (d) – (f) Relationships between discrimination and finger properties, with highest correlation for finger stiffness, then finger size, and ridge breadth.

The finger properties of each participant, including finger size, skin stiffness, and fingerprint ridge breadth, were measured, as detailed in Methods, with variance observed across the cohort. As indicated in Fig. 1(b), the male participants of this study spanned a range of finger widths of more than one standard deviation (mean = 18 mm) as compared with the literature (Tilley & Associates, 2001), while those for the female participants were closer to the average value (mean = 15 mm) within one standard deviation. Within our study population, a high correlation was observed between finger size and skin stiffness (*R* = 0.90, p-value < 0.001, SE = 2.96 x 10^-4^), with a less significant correlation between either of these metrics and fingerprint ridge breadth, Fig. 1(c). Fingerprint ridge breadth was correlated with finger size (*R* = 0.76, p-value < 0.001, SE = 9 x 10^-4^), as noted elsewhere (Loesch & Martin, 1984*a*; Dillon *et al*., 2001), and with skin stiffness (*R* = 0.70, p < 0.001, SE = 4.4 x 10^-3^).

While all three finger properties correlated to some extent with perceptual performance in the compliance discrimination task, skin stiffness exhibited the highest correlation across the stimulus pairs (*R* = -0.95, -0.93, and -0.77 for 184/121, 33/5, and 75/45 kPa pairs), followed by finger size (*R* = -0.91, -0.82, and -0.65) and ridge breadth (*R* = -0.81, -0.70, and -0.43), respectively, Fig. 1(d – f). Together, these results indicate that although finger size and skin stiffness are correlated, skin stiffness tends to be more strongly correlated with perceptual performance.

### Experiment 2: Spatiotemporal characterization of skin surface deformation

Local patterns in the deformation of the skin surface constitute the major factor driving our percept of compliance (Srinivasan & LaMotte, 1995; Ambrosi *et al*., 1999; Moscatelli *et al*., 2016; Dhong *et al*., 2019; Li *et al*., 2023), although proprioceptive finger movements can provide supplementary information (Xu *et al*., 2021*a*). To evaluate if skin deformation cues align with trends in both perceptual discrimination and finger property measurements, we imaged 3-D high-resolution profiles of the skin’s surface, using techniques described previously (Hauser & Gerling, 2018*a*; Li *et al*., 2020; Li & Gerling, 2021; Li *et al*., 2023), Fig. 2. Basically, images taken by two cameras through transparent, compliant substrates are used to construct 3-D point clouds which represent the geometry of finger pad at the contact surface. This is done for the seven compliant stimuli ranging from 5 to 184 kPa. From the point cloud data, we evaluate five derived cues to characterize the skin surface’s deformation over time, which include contact area, curvature, penetration depth, eccentricity, and force (Li *et al*., 2020).

**Figure 2.**
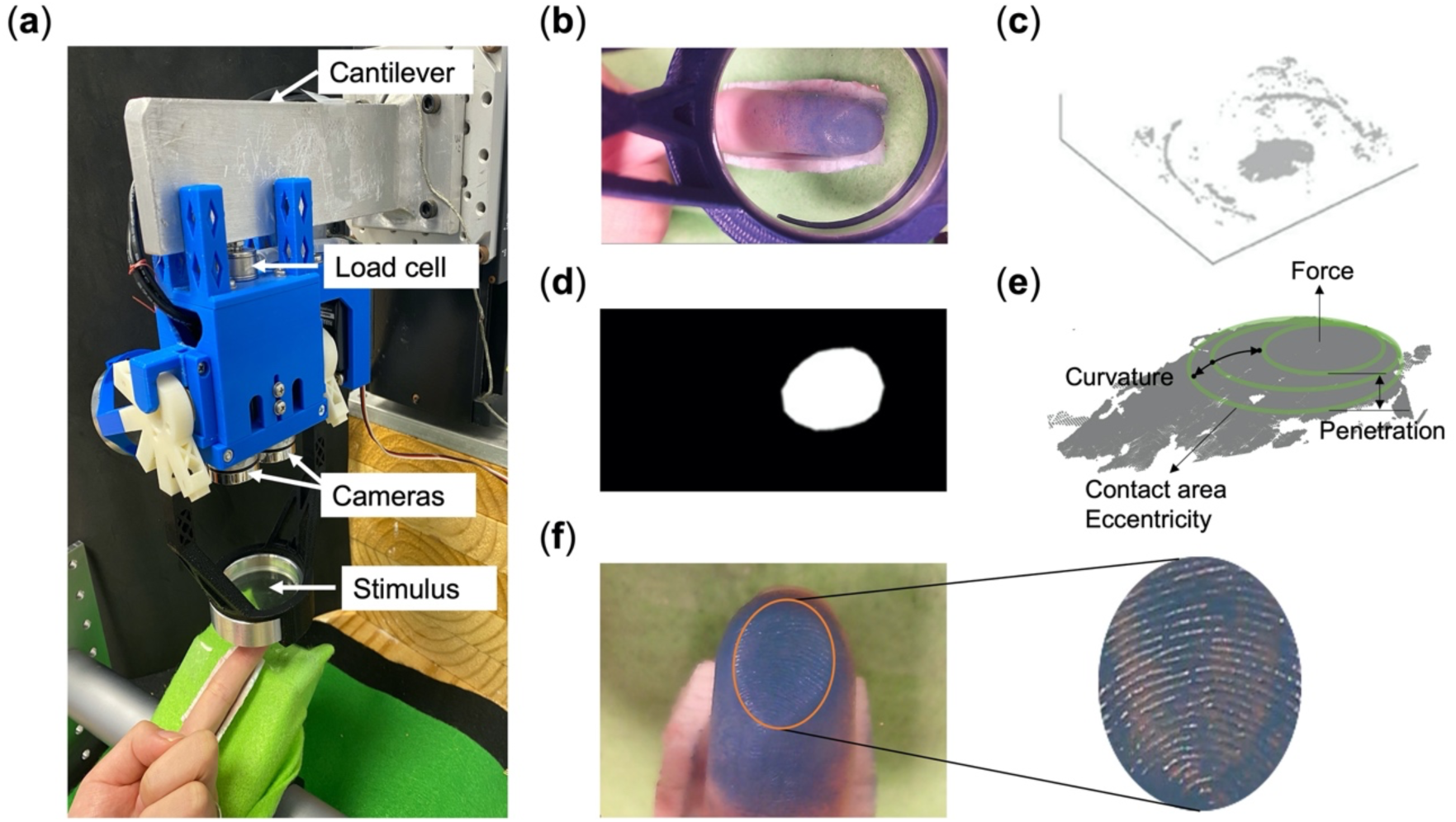
Experimental setup to generate 3-D point clouds for evaluating finger skin deformation. (a) Mechanical-electrical indenter delivers elastic, compliant substrates to fingertip. (b) Left camera image of fingertip indented by 45 kPa substrate at 2 mm displacement. (c) 3-D point cloud is constructed from left and right images of the finger using a disparity-mapping algorithm. (d) Region of contact between skin and substrate is manually selected by masking. (e) Clean point cloud that represents the skin surface is generated, ellipses are fitted to reduce its dimensionality, and five skin cues quantify patterns of skin deformation. (f) Glass plate is used in separate experiments to measure each participant’s skin stiffness and ridge breadth.

From representative point cloud data in Fig. 3, one can observe that the skin deforms distinctly with respect to stimulus compliance and skin stiffness. Indented by a less compliant stimulus (184 kPa, row 3 in Fig. 3(a)), the finger pad’s surface flattens against the substrate, which generates higher contact area and force with a more circular contact shape, i.e., lower eccentricity (blue lines in Fig. 3(b)). In contrast, with a more compliant stimulus (45 kPa, row 1 in Fig. 3(a)), the finger pad’s surface retains its shape and penetrates into the bulk of the substrate, which generates greater curvature and penetration depth with a more elliptical contact shape, i.e., higher eccentricity (green lines in Fig. 3(b)). Moreover, an individual’s skin stiffness also affects how the skin surface deforms. For instance, when the more compliant stimulus (45 kPa) is indented into softer skin (0.097 N/mm), we observe less contact area as opposed to stiffer skin (0.152 N/mm) (rows 1 and 2 in Fig. 3(a) and two green lines in Fig. 3(b)). However, the differences in contact area between these participants are less pronounced than those between the stimulus compliances (45 and 184 kPa). The distinctions between participants, as observed for contact area, do not follow for certain other cues, such as for force, as with the more compliant 45 kPa stimulus. This figure presents an illustrative example, in the subsequent section the data for all participants are statistically evaluated.

**Figure 3.**
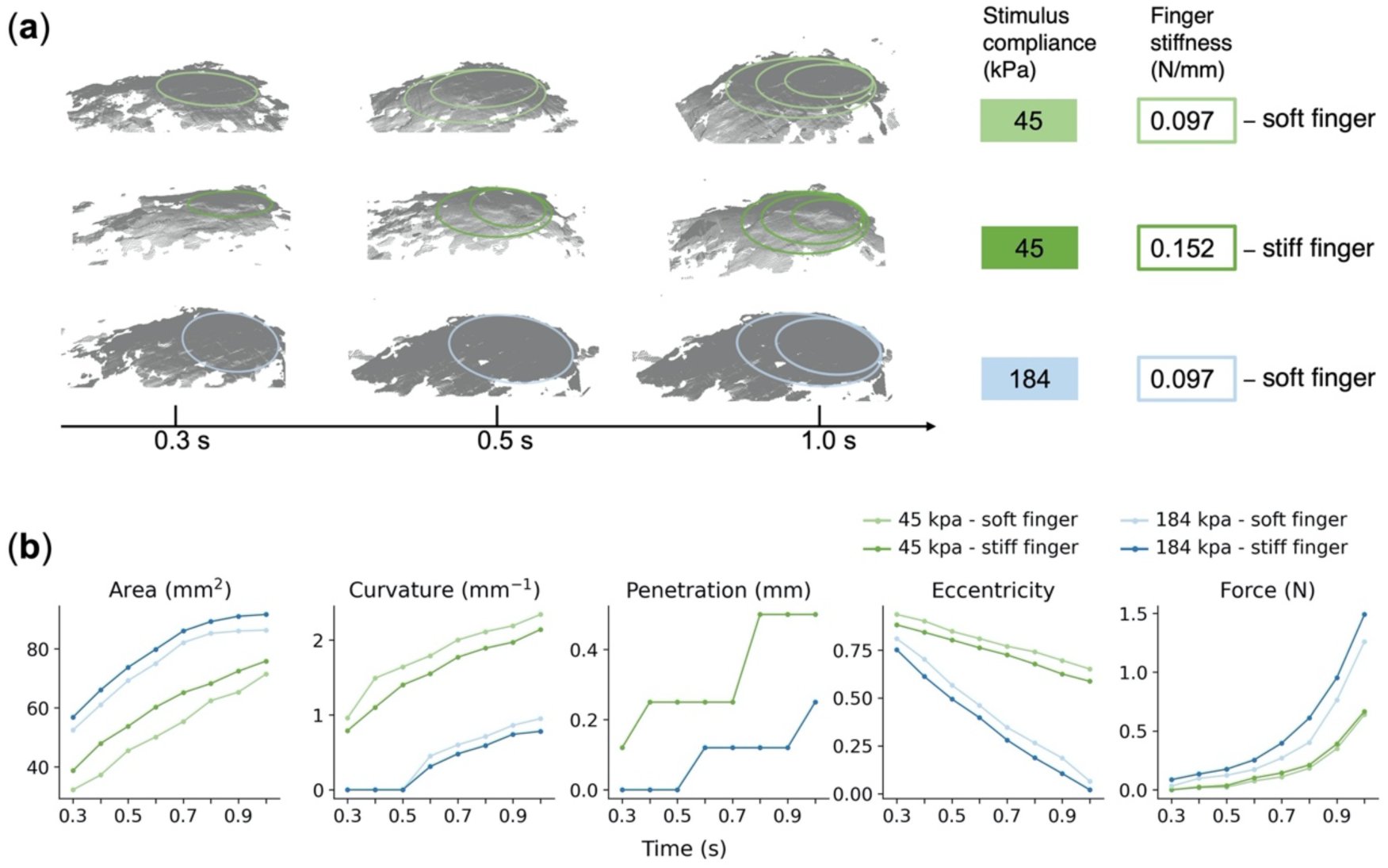
Time-evolution of 3-D skin deformation for stiff and soft fingertips indented by stiff and soft stimuli. (a) 3-D reconstructions of skin surface deformation at 0.3, 0.5, and 1 sec time-points into the 2 mm indentation at a rate of 1.75 mm/sec. Stiff (0.152 N/mm) and soft (0.097 N/mm) fingertips contact more (45 kPa) and less (184 kPa) compliant stimuli. (b) Five skin deformation cues between the two participants and two stimulus compliances. Data points represent means of three trials per participant. Contact area as well as other cues differ most notably between the two compliances, and to a relatively lesser, but still prominent extent between participants. These example cases are subsequently quantified and statistically evaluated across a population of participants, Fig. 4.

### Experiment 3: Individual differences in finger properties modulate skin deformation and perceptual discrimination

Though perception is most highly correlated with skin stiffness, as shown in Fig. 1(d), we next question if differences in participants’ skin stiffness evoke patterns of skin deformation associated with improved perceptual discrimination. To address this question, in Fig. 4(a-b) the rate of change in skin deformation per participant is compared, between compliant pairs, with the corresponding rate of correct discrimination. The rate of change, as opposed to a static value or a terminal value, is used as temporal changes in contact area and force have been found to be most predictive of perceptual discrimination of compliance (Srinivasan & LaMotte, 1995; Ambrosi *et al*., 1999; Moscatelli *et al*., 2016; Dhong *et al*., 2019; Xu & Gerling, 2020; Xu *et al*., 2021*a*; Li *et al*., 2023). Here, we define the change rate in contact area as the median of the discrete intervals calculated per 0.1 s, as indicated by the denoted line segments in Fig. 4(a) for the 184 and 121 kPa stimuli, which exhibit median change rates of 45 and 50 mm^2^/s in contact area, respectively. Indeed, the results of Fig. 4(b) also show that two of the cues, i.e., change rates of contact area and eccentricity, are highly correlated with perceptual discrimination across the cohort of 25 participants in distinguishing the 33/5 and 184/121 kPa stimulus pairs. With these two stimuli, we observe high correlation values of 0.9 and 0.87 with contact area, and of 0.86 to 0.84 with eccentricity. In contrast, the change rates of force and curvature are correlated with perceptual discrimination for only the less compliant (184/121 and 75/45 kPa) and more compliant (33/5 kPa) pairs, respectively. Similar trends have been observed previously, with regard to force cues, which are predictive of perceptual acuity when the stimuli are stiffer than the skin (Friedman *et al*., 2008*a*; Kaim & Drewing, 2011; Hauser & Gerling, 2018*b*; Li *et al*., 2020).

**Figure 4.**
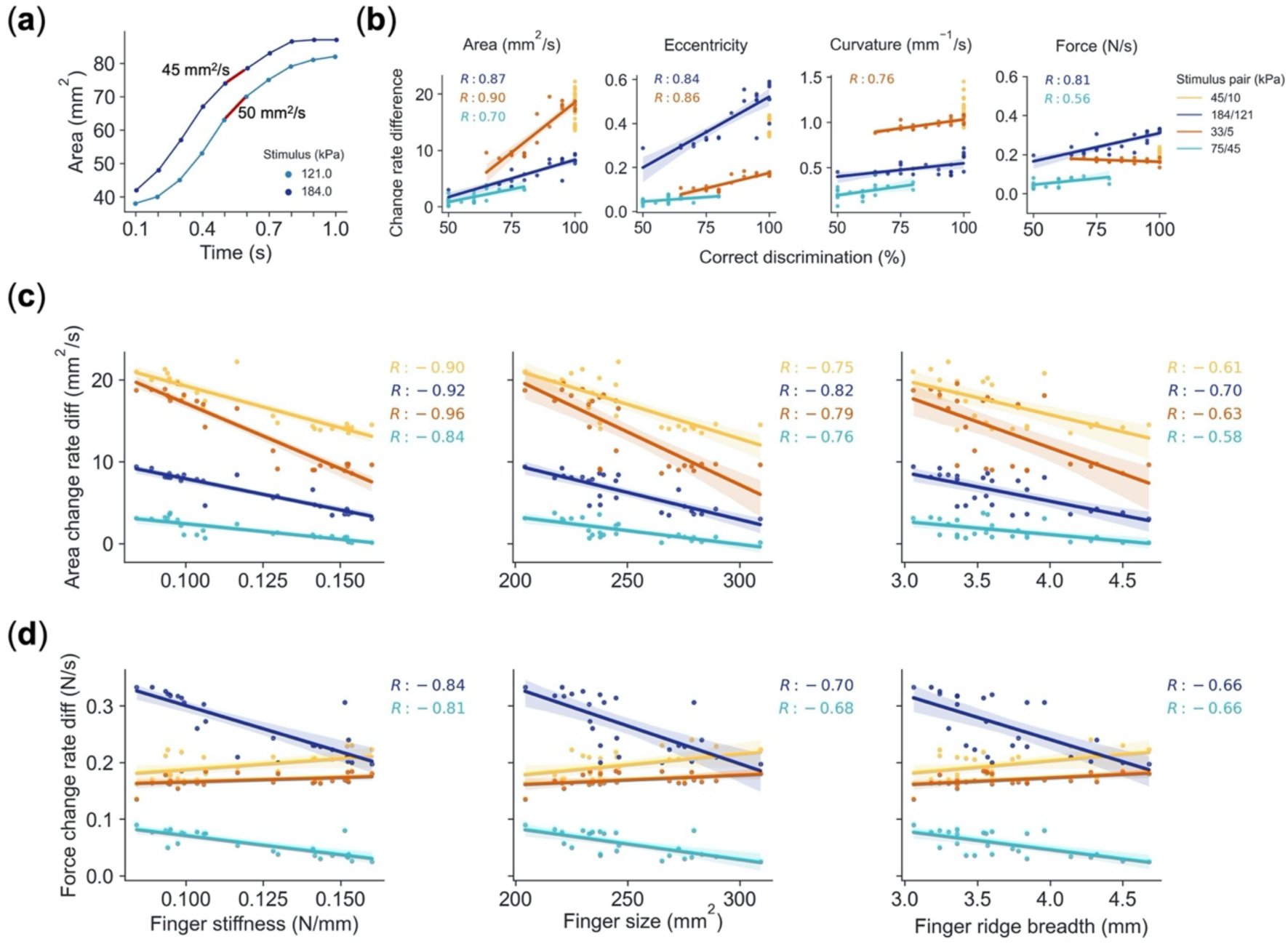
Relationships between skin deformation cues and perceptual discrimination, and finger properties of stiffness, size, and ridge breadth. (a) Steps in calculating change rate in contact area every 0.1 sec, for 184 and 121 kPa stimuli, with median values of 45 and 50 mm^2^/s. Completed across all trials, one data point is plotted per participant in subsequent panel. (b) Change rates in contact area of 25 participants are highly correlated with discrimination across 184/121, 33/5, and 75/45 kPa compliant pairs. Change rate in force correlates with discrimination for stiffer 184/121 stimuli. (c) Finger stiffness correlates with change rate in contact area, followed by size and ridge breadth. (d) Finger stiffness also highly correlates with change rate in force, but only for less compliant pairs.

Next, in Fig. 4(c, d) we evaluate how finger stiffness, size, and ridge breadth influence the change rate of contact area and force. Across either skin deformation cue, skin stiffness is the most highly correlated among the finger properties. Indeed, the change rate of contact area is highly correlated with skin stiffness (*R* > 0.9) across 45/10, 184/121, and 33/5 kPa pairs, in contrast to finger size (*R* = 0.75, 0.82, and 0.79). Moreover, those participants with softer finger stiffness generate the largest distinctions between the stimulus pairs, as is observable in the line slopes. Similarly, the change rate of force shows significant correlation with skin stiffness, but only for the less compliant 184/121 and 75/45 kPa pairs (*R* > 0.8), where those participants with softer fingers generate the largest differences, Fig. 4(d).

### Experiment 4: Decoupling the relative impacts of finger stiffness and size on perceptual discrimination in subgroups of participants

We have shown finger size and skin stiffness are highly correlated in Fig. 1(c) and both contribute to perceptual acuity by tuning patterns of skin deformation at the point of contact surface, Fig. 4(c, d). Therefore, we developed statistical models to decouple the relative contributions of these finger properties. Using a K-means approach to form clusters of participants, based on the three finger properties (stiffness, size, and ridge breadth), we identify two distinct groups of participants, Fig. 5(a) and Appendix A3. Even though the groups mostly separate upon lines of gender, a few male participants with small or soft fingers are categorized in Group 1, which contains participants with smaller and softer fingers, including all of the female participants. Perceptual discrimination between these two subgroups is found to be significantly different for the 184/121 kPa pair (t-statistic = 5.1, p < 0.001), the 33/5 kPa pair (t- statistic = 4.2, p < 0.001), and the 75/45 kPa pair (t-statistic = 2.35, p = 0.03), which is not discriminable above the 75% threshold for either Group 1 or 2, Fig. 5(b). Only Group 1’s performance exceeds a 75% threshold for the 184/121 and 33/5 kPa compliant pairs. Moreover in Fig. 5(c), principal component analysis (PCA) on the skin deformation cues shows that the skin deforms distinctly between the two groups, with the small and soft fingers of Group 1 more pronounced in contact area, penetration depth, and curvature, whereas the larger and stiffer fingers of Group 2 are more pronounced in force and eccentricity. This trend aligns with the finding above that a nominal finger flattens and exhibits less 3D curvature against a stiffer substrate, Fig. 3(b, curvature) and indeed in this regard stiffer finger skin is equivalent to a stiffer substrate.

**Figure 5.**
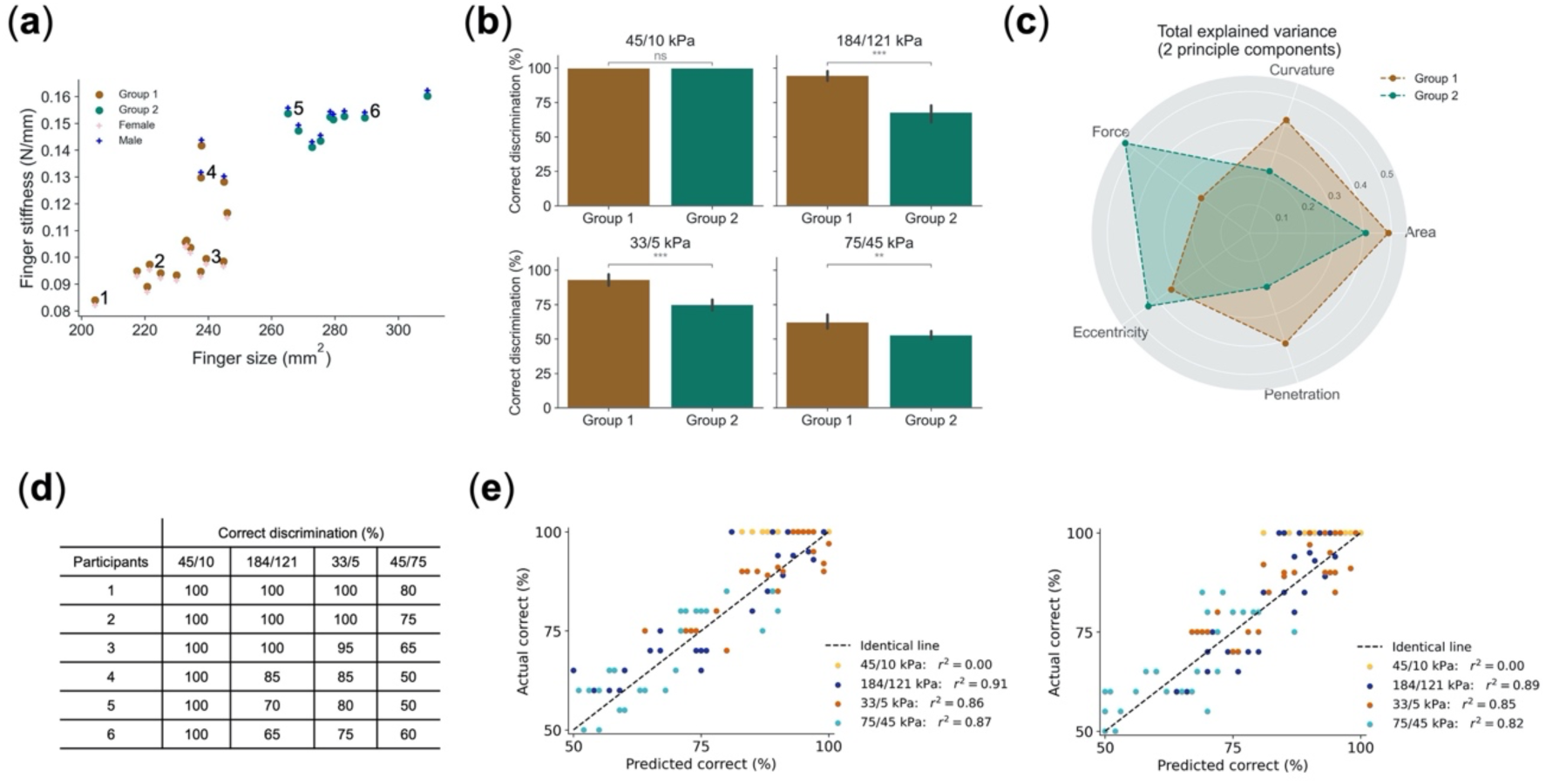
Multivariant regression models separate distinct cohorts, with discrimination predicted by finger stiffness. (a) Based on finger size, stiffness, and ridge breadth, K-means clustering splits participants into two cohorts, mostly of male and female participants, with some males with smaller and softer fingers in Group 1. (b) Discrimination differs between groups (***p<0.001, **p<0.05). (c) Utility of skin deformation cues between groups differs, e.g., Group 2 with stiffer skin relies on force cues. (d) Discrimination of six participants from panel (a) differs based on finger stiffness and size. (e) Left: Evaluation of regression model where perceptual discrimination is predicted by finger size, stiffness, and ridge breadth. Right: Similar prediction with finger stiffness alone. Negligible decrease in performance lends support to prominent role of skin stiffness.

To gain an intuitive understanding of the relative effects of finger size and stiffness, the psychophysical performance of six select participants, as indicated by the participant numbers in Fig. 5(a), was then compared in Fig. 5(d). As expected, the highest discrimination performance is observed for participant 1 with the smallest and softest finger. In contrast, when finger size is similar but skin stiffness varies (participants 3 and 4 from Group 1), lower skin stiffness (participant 3) leads to higher discriminative performance. In contrast, when the skin stiffness is similar but finger size varies (participants 2 and 3 from Group 1 and participants 5 and 6 from Group 2), a large increase in finger size does not necessarily result in better performance. This case-by-case observation aligns with Experiment 3, Fig. 4(c) and suggests that skin stiffness affects perceptual discrimination more than finger size.

More quantitatively, multivariant linear regression models were built to predict individuals’ perceptual discrimination in two conditions. The first was built to predict a participant’s psychophysical performance from all three skin properties of finger size, stiffness, and ridge breadth, while the second was built based upon skin stiffness alone. Comparing the predictive accuracy of these models, the results indicate only slight improvements are obtained upon adding the factors of size and ridge breadth, Fig. 5(e), which further lends support to the prominent role of skin stiffness over the other finger properties.

### Experiment 5: Directly modulating skin stiffness improves one’s perceptual discrimination

As a more direct means of determining the role of skin stiffness in discrimination, we used hyaluronic acid, a common moisturizer, to directly soften the skin stiffness of a new cohort of 15 participants. In specific, a drop of hyaluronic acid was applied to the index finger, which was not washed immediately prior, and allowed 1-2 minutes to dry. The results show this treatment significantly and systematically reduces participants’ skin stiffness, Fig. 6(a). As well, upon applying hyaluronic acid, participants’ discrimination performance also significantly improves, especially for those with relatively stiffer fingers (participants 9-15), Fig. 6(b). Upon using clustering to separate the cohort of 15 participants into two distinct groups, we find that patterns of skin deformation, especially the change rate of contact area increases significantly across the three stimulus pairs for Group 2, Fig. 6(c). We also find that the discriminative performance between the groups improves for the 75/45 kPa pair in Group 1 (t-statistic = -13.3, p < 0.001), which is the only case where this group had performed below a 75% threshold without hyaluronic acid, and for all three pairs in Group 2 (t-statistic = -5.9, p < 0.001 for 184/121 kPa, t-statistic = -5.7, p < 0.001 for 33/5 kPa, t-statistic = -7.8, p < 0.001 for 75/45 kPa pairs), Fig. 6(d). The results suggest that direct measures to soften skin stiffness improve perceptual performance, at both individual and aggregated levels.

**Figure 6.**
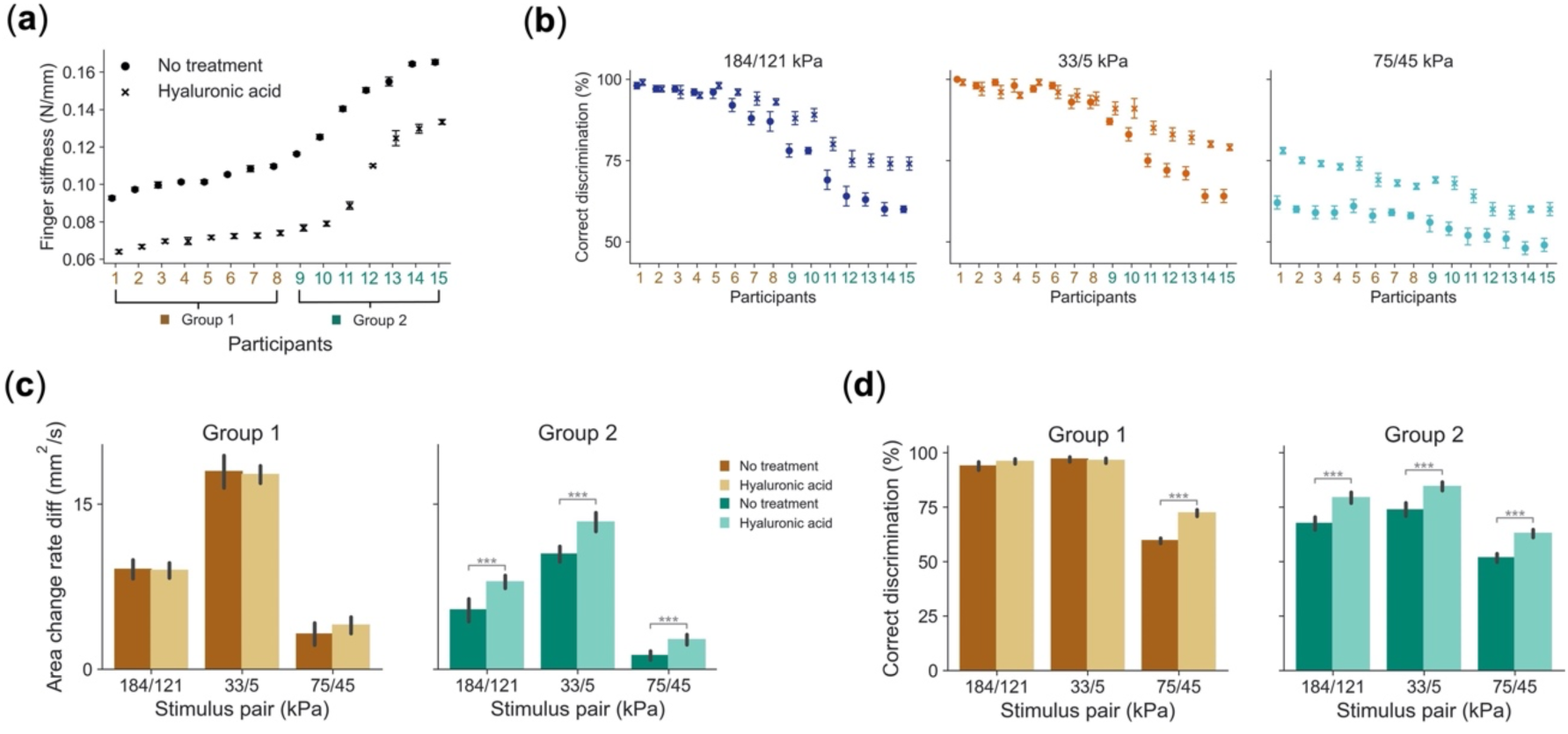
Skin stiffness is modulated by hyaluronic acid, which systematically increases change rate in contact area and improves discrimination. (a) Skin stiffness before and after applying hyaluronic acid. (b) Discrimination of compliant pairs, before and after applying hyaluronic acid, where performance improves. (c) Distinct cohorts are separated by K-means clustering, based on finger stiffness before skin treatment. Patterns of skin deformation are significantly accentuated for Group 2 participants (***p<0.001). (d) Discrimination also improves for Group 2 participants, who performed worse than Group 1 without treatment (***p< 0.001). Note that Group 1 achieves nearly perfect discrimination in 184/128 and 33/5 kPa cases, even without treatment, and therefore sees little improvement. However, with 75/45 kPa stimuli, which are more difficult to discriminate, both groups improve.

## DISCUSSION

While declines in tactile acuity with aging are a function of multiple factors, for younger people the current working hypothesis has been that smaller fingers are better at informing perceptual discrimination due to a higher density of neural afferents. In helping to refine this theory, the studies herein show that an individual’s perceptual acuity – while highly coupled with finger size – is more robustly correlated with and predicted by their skin stiffness. As shown via imaging of the skin surface, softer skin deforms at a higher rate at the point of contact. These higher rates of change correlate with participants’ abilities in perceptual discrimination of compliant pairs of stimuli. It is likely that higher rates of change in surface contact govern the temporal recruitment of spatial populations of cutaneous afferents. Moreover, we show that each individual’s skin can be softened, upon the application of hyaluronic acid, to generate higher rates of change in surface contact area and immediate improvements in perceptual acuity. Overall, these findings shed light on the role of the skin’s elasticity in shaping and modulating the encoding of touch stimuli and the bases of individual differences in acuity. Indeed, although influenced by one’s finger size, and geometry at the point of contact, an individual’s level of tactile acuity can be significantly improved by modulating their skin stiffness.

### Role of skin stiffness in shaping individual differences in perception

The skin’s degree of elasticity enhances local patterns of skin deformation when contacting objects. We performed an extensive set of experiments in attempt to evaluate the interplay between skin stiffness and finger size, based upon methods to image the dynamics of the skin’s surface deformation. The imaging shows that deformation patterns are highly correlated with perceptual acuity, notably the rate of change in contact area, Fig. 4(b), in alignment with prior reports (Srinivasan & LaMotte, 1995; Ambrosi *et al*., 1999; Moscatelli *et al*., 2016; Dhong *et al*., 2019; Xu & Gerling, 2020; Xu *et al*., 2021*a*; Li *et al*., 2023). Then, we observe that softer fingers generate larger rates of change in contact area between pairs of stimuli, Figs. 4(c), 6(c), as compared to either finger size or ridge breadth. A lower elastic modulus of the skin has been shown in elastography to affect the distribution of stresses and strains throughout its layers (Yang *et al*., 2018) and in neural computational simulation to improve the encoding of stimuli (Deflorio *et al*., 2023). Such distinctions at the skin surface are thereby likely propagate to locations of mechanosensitive end organs at the epidermal-dermal border and influence neural firing. To further evaluate the influence of the skin’s stiffness on perceptual outcomes, this time in a direct fashion within individuals, we applied hyaluronic acid. By applying hyaluronic acid at the skin surface, the initially observed range of stiffness from 0.09 to 0.17 N/mm across the participants, Fig. 6(a), was systematically reduced by about 40%, and correspondingly, perceptual discriminability improved by 5% to 30%, Fig. 6(b, d). The discriminative ability of each participant immediately improved, especially for those with the stiffest skin. Indeed, similar to observations between participants in Fig. 4(c), we show within participants that one’s rates of change in contact area increases when his or her skin is softened, Fig. 6(c). Indeed, more elastic skin generates higher rates of change in contact area during initial contact with an object.

Our finding of the direct impact of skin mechanics on tactile acuity aligns with those of prior neurophysiological and perceptual studies, as well as computational models. For instance, others have shown that skin properties influence neural encoding, in both the responsiveness of single afferents and the recruitment of populations of afferents. Regarding the former, Hudson, et al. (2015) directly modulated the compliance of the finger pad, using venous occlusion in placing an inflated cuff around the arm resulting in engorgement of the fingers. This intervention significantly influenced the firing rates of subtypes of cutaneous mechanoreceptors, especially rapidly adapting afferents (Hudson *et al*., 2015). Likewise, in the realm of computational modeling, finite element models have shown that increases to the simulated skin’s stiffness generate higher concentrations of stresses and strains near interior locations of mechanosensitive end organs (Gerling & Thomas, 2008; Gerling, 2010; Pham *et al*., 2017; Hamasaki *et al*., 2018). Regarding the response of a populations of afferents, force-related cues have been correlated with the total number of active fibers (Friedman *et al*., 2008*b*; Saal *et al*., 2009), and the change rate of contact area is likely better related to the number and rate of afferents recruited, though not measured in that study. Emerging efforts to record afferent population responses using calcium imaging in mice may open new avenues for a direct coupling with patterns of skin surface deformation (Broussard *et al*., 2018; Moayedi *et al*., 2023).

Others have observed that upon hydrating the skin its stiffness increases (Sandford *et al*., 2013), as well as its surface contact area (Serhat *et al*., 2022), which aligns with our finding that softer fingers generate higher rates of skin deformation. Moreover, skin’s stiffness can also be internally regulated. In particular, sweat ducts very densely populate the fingerprint ridges to moisturize and soften to the skin to help regulate one’s grip to avoid severe object slips (André *et al*., 2010; Yum *et al*., 2020). In this study, we used compliant pairs of stimuli. As well, even with other stimuli, e.g., rigid gap stimuli to evaluate neurological deficits, those participants with higher skin conformance achieve lower detection thresholds (Woodward, 1993; Vega-Bermudez & Johnson, 2004; Gibson & Craig, 2006). Overall, in extending the prior studies, our results further indicate the elasticity of the skin helps evoke changes in rates of change of contact area that may lead to the incremental rate by which neural afferents are recruited, which may be key to encoding discriminable differences.

### Decoupling influences of skin stiffness, finger size, and gender

Despite the prominent role of skin stiffness in modulating patterns of local skin deformation, it is clear that finger size is also an important factor in governing tactile acuity. Indeed, across the participants, we observed that finger size and skin stiffness were highly correlated (*R=0.90*), Fig. 1(c). Moreover, the highest performance tends to be observed for those with the smallest fingers, Fig. 5(a,b), even when participants’ level of acuity is modulated with hyaluronic acid. In Fig. 6(a), while we observe that the highest values of untreated finger stiffness are reduced by applying hyaluronic acid, their skin stiffness is not reduced to levels observed for those with the softest untreated fingers. For example, participant 15’s treated finger skin stiffness is reduced to 0.13 N/mm, but this does not reach participant 8’s untreated value of 0.11 N/mm. Where similar levels of skin stiffness are achieved, e.g., participant 11, the improvement in perceptual discrimination is observed from 69 to 80% for the 184/121 kPa compliant pair, which is not quite to the level of 87% observed for participant 8 in the untreated skin case. Therefore, though it is clear that skin stiffness can be modulated, and such modulation does increase the rate of change in contact area, and thereby tactile acuity, the size of one’s finger does eventually limit performance. Therefore, with the same number of afferents between people, and similar finger stiffness, it appears that a smaller finger will exhibit a higher density, and therefore smaller increments in the change in contact area will impact the rate of afferents recruited. Also of note, prior studies have shown that perceptual acuity tends to play out along gender lines. In particular, women exhibit higher acuity and on average have smaller fingers with higher receptor densities (Woodward, 1993; Peters *et al*., 2009; Abdouni *et al*., 2018, n.d.). In this study we found that individuals with small fingers also tend to have soft fingers, Fig. 1(c), tend to be women, Fig. 5(a), and outperform individuals with larger and stiffer fingers in discrimination tasks, Fig. 1(d, e). These findings therefore replicate those prior with respect to finger size and gender. However, we did observe cases of men with smaller and softer fingers, Fig. 5(a, d), whose performance nearly matched that of the cohort of women with improved levels of perceptual acuity.

### Larger fingers do not generate higher contact area

We analyze the variable of rate of change in skin deformation, particularly contact area, between compliant stimuli and the skin surface during the course of initial contact. Given prior literature on finger size and innervation density, one might assume that larger fingers generate larger absolute contact area, in general, and therefore if greater contact area might recruit more afferents, or at a larger rate. In Appendix A5, we compare contact area measurements across participants at a consistent time point of 0.3 sec of indentation into a glass plate, a time point that is selected because it occurs before 0.4 sec when stimuli start to become reliably discriminable (Li *et al*., 2023). From Appendix A5(a, b), we observe that even for those participants with larger fingers, their contact areas are not systemically related to their finger sizes, nor does this hold for skin stiffness. The lack of relationship is likely due to individual differences in geometry at the tip of the finger (Harih & Tada, 2015; Voss & Srinivasan, n.d.), which we did not measure. In particular, we measured finger size at the distal phalange, to obtain greater consistency among participants, while contact area imaging measurements were made at the fingertip, with it positioned at a 30 degree angle. Additionally, the lack of relationship for skin stiffness indicates that the initial contact area does not significantly impact measurements of skin stiffness.

Moreover, in Appendix A5(c), we see that a participant exhibiting greater contact area does not correlate with perceptual performance. On one hand, these data lead us back to the importance of rate changes in skin deformation, Fig. 5(c). Perhaps more importantly, they indicate that amidst randomness and variability in finger geometry, the governing of tactile acuity by skin stiffness, as opposed finger size, is an effective way to neutralize potential impacts on perception. Indeed a population of people can exhibit larger variance in finger size and finger geometry, yet fingers stiffness can help overcome these.

### Adjusting active touch strategies amidst changes in skin stiffness

This study was conducted according to a passive touch paradigm, and we observed immediate improvements in each individual’s tactile acuity upon the softening of their skin. Such changes in the skin’s mechanics may also influence our exploration strategies and behaviors in more natural situations of active touch. Indeed, in active touch, certain exploration strategies may improve discriminative performance. For example, those with stiffer fingers tend to apply higher forces and larger displacements when differentiating compliant objects (Xu *et al*., 2021*b*). Additional work is needed to better understand how softening the skin might impact an individual’s deployment of active touch strategies. We have reported elsewhere that compliant pairs can be discriminated at 0.4 sec, depending on stimulus elasticity and velocity (Li *et al*., 2023), which is very early into the movement of the finger. In passive touch, we controlled the velocity ramp between the pair of stimuli. This was done because when stimulus velocity is controlled in passive touch, what differs between the stimulus pair is the change in force rate. The rationale for this approach is that it that affords perceptual judgments based on the most minimal levels of skin deformation, especially with stimuli less compliant than skin (Bergmann Tiest & Kappers, 2009; Hauser & Gerling, 2018*b*), which conserves energy, and is tied with human volitional touch strategies (Xu & Gerling, 2020; Xu *et al*., 2021*a*, 2021*b*).

## MATERIALS AND METHODS

### Participants

A total of 40 young participants were recruited (mean age = 28, range = 21 – 32; 19 males and 21 females). Of these, a cohort of 25 participated in Experiments 1-4, with a separate cohort of 15 in Experiment 5. None had prior experience with our experimental tasks or reported a history of finger injury. Their fingers were free of calluses and scars. A written, informed consent was obtained from each participant before conducting the experiments, which were approved by the local institutional review board. All devices and surfaces were sanitized regularly. All participants wore facemasks during the experiments, following COVID-19 protocols.

### Stimulus fabrication

As a point of reference, the average compliance of a human finger skin is about 42 – 54 kPa (Miguel *et al*., 2015; Oprişan *et al*., 2016). Seven elastic stimuli were fabricated with compliance values ranging from 5 to 184 kPa, in which one stimulus (45 kPa) was close in stiffness to that of human skin, with three softer (5, 10, 33 kPa) and three harder (75, 121, 184 kPa). Each silicone-elastomer formulation was poured into an aluminum collar (5.4 cm outer radius by 1.6 cm depth) fitted and sealed with a clean, dry glass disc (5.1 cm radius by 0.3 cm depth) using a syringe tip filled with 0% diluted silicone rubber, heated at 100 C until fully sealed. To formulate the more compliant stimuli (5, 10, 33 kPa), a two-component silicone rubber (Solaris, Smooth-on Inc., Macungie, PA, USA) was mixed at ratio of 1:1, diluted by silicone oil (ALPA-OIL-50, Silicone oil V50, Modulor, Berlin, Germany) with ratios of 200 wt.% for 33 kPa, 300 wt.% for 10 kPa and 400 wt.% for 5 kPa. Each mixture was placed in a vacuum chamber under 29 mmHg for 5 min and then allowed to rest at room temperature until all air bubbles were released. Next the transparent mixture was cured in an oven at 100 C for 25 min to fully solidify before returning to room temperature. To eliminate surface stickiness caused by high dilution ratios, a thin layer of 100% diluted silicone rubber was applied on the stimulus’s surface, then cured at 100 C for 15 min. For the rest of stimuli (45, 75, 121, 184 kPa), a different two-component silicone rubber (Sylgard 184, Dow Corning, Midland, MI, USA) was used and diluted by silicone oil with ratios of 200 wt.%, 100 wt.%, 50 wt.% and 0 wt.%, respectively.

### Measurements of finger skin properties

#### Finger size

Measurements by digital caliper at an interval of 0.1 mm of the length and width of distal phalanx of the index finger were used to evaluate participants’ finger size. The length is measured as the distance between finger edge and DIP joint, and the width is the distance from left to right edge of the finger, where they intersect the center of finger pad. From these measurements, finger size is estimated as the area of the ellipse formed by the length as its major axis and width as the minor axis.

#### Finger stiffness

Finger stiffness is typically measured by a rigid body compression test (Hyun- Yong Han & Kawamura, 1999). A glass plate (5.4 cm radius by 0.3 cm depth) was indented in the index finger at a rate of 1 mm/s, while simultaneously measuring force. The stiffness is approximated as the slope of linear regression of the force-displacement curve, Appendix A1.

Though stiffness measurements with the literature can vary due to the exact experimental setup, under a plate displacement of 2 mm and contact angle of 30 degree, our force readings (1.87 N) align with the values of 2 N and 1.9 N reported in the literature, which used rigid plates (Hyun- Yong Han & Kawamura, 1999) and hard elastic stimuli (Hauser & Gerling, 2018*b*), respectively. Moreover, since initial contact area, determined by finger size, has potential influence on force, we evaluate the correlation between contact area and stiffness measurements within the linear region of force-displacement curve, Appendix A5(a). The results indicate that initial contact area did not have a significant impact on the measurements of finger stiffness (*R* = 0.27).

#### Finger ridge breadth

During its indentation by a glass plate, images of finger contact were captured by the indenter cameras and the contact surface between skin and glass plate was extracted, Fig. 1(b) and Appendix A2. To obtain a clear representation of the fingerprint ridge lines, we applied a localized thresholding filter (OpenCV, Python 2.8) with adjusted contrast and brightness in the grayscale image, Fig. 1(b). Then, measurements of ridge breadth were acquired using a published approach (Mundorff *et al*., 2014), where the ridge breadth was measured as the length of a line crossing ten parallel ridges with no minutiae causing a break in the perpendicular line. Furthermore, we converted the units from pixels to millimeters, Appendix A2(b).

### Apparatus

The experimental apparatus utilizes a custom-built, mechanical-electrical indenter with a stereo imaging system, Fig. 2(a). Full details of its setup and validation have been previously published (Hauser & Gerling, 2018*a*; Li *et al*., 2020). Briefly, the apparatus consists of a motion-controlled indenter and load sled (ILS-100 MVTP, Newport, Irvine, CA, USA), a load cell on top of a customized stimulus housing, and two stereo cameras oriented in parallel. Up to five substrates can be delivered individually to the finger pad by rotating the stimulus housing, at controlled delivery rate, displacement, and duration. A support platform was designed to stabilize the participant’s hand with a solid plastic housing to minimize finger movement during the experiment. The finger was adjusted to 30-degree angle with respect to the stimulus surface.

Indentation displacement was recorded by the motion controller and the force was measured by the load cell at 150 Hz, with a resolution of ± 0.05 N (LCFD-5, Omegadyne, Sunbury, OH, USA). Th two stereo webcams (Papalook PA150, Shenzhen Aoni Electronic Industry Co., Guangdong, China) installed vertically above the stimulus captured images at 30 frames per second, with a maximum resolution of 1280 by 720 pixels.

### Image acquisition and processing methods

#### Generating 3-D point clouds

The procedures of reconstructing 3-D surface of finger pad are illustrated in Fig. 2(b, e), and published previously (Hauser & Gerling, 2018*a*). A thin layer of ink dots were applied on the finger using a paint brush with its bristles normal to the skin surface. Then, images were taken by the left and right cameras simultaneously during indentation. A disparity-mapping approach was used to co-locate the ink points on the skin surface between the two images, as a result, a point cloud was generated using the identified pixel brightness values as the coordinates in the 3-D domain. Then we extracted the region of contact between the skin and substrate by masking the remaining areas. Finally, a clean 3-D point cloud that represents the skin surface of finger pad was constructed. On average, each point cloud has 30,000 discrete points after imaging processing. A unique point cloud was generated every 100 ms to provide high spatiotemporal resolution for analyzing skin deformation.

#### Ellipse method and derived skin deformation cues

To reduce the dimensional complexity but sufficiently capture the characteristics of the skin’s deformation, we developed a method that fits vertically stacked ellipses with same orientation onto the point clouds (Li *et al*., 2020). The fitting process starts from the contact surface to the highest penetration of the finger pad, i.e., the bottom ellipse to the top in Fig. 2(e). Each ellipse is defined as an image plane in which the plane of contact surface is defined as the first image plane. Each image plane contains at least 98% of the points with 95% confidence and the distance between two adjacent image planes is 0.25 mm, which is twice the resolution of the stereo images in the vertical dimension (0.12 mm). As a result, a point cloud containing nearly 30,000 discrete points is converted into about 1-6 ellipses. Next, using this ellipse’s representation, we characterize the dynamic change of skin by developing five skin deformation cues, including contact area, curvature, penetration depth, eccentricity and force. The algorithmic definition of each cue has been published previously (Li *et al*., 2020).

### Experimental design

#### Psychophysical experiments

A series of discrimination tasks were designed and conducted to evaluate the participants’ discrimination ability in compliance. Four pairs of stimulus compliances were prepared as 45/10, 184/121, 33/5 and 75/45 kPa, where 184/121 kPa pair is less compliant than that of the skin, and 33/5 kPa pair is more compliant than the skin, and the remaining pairs span the skin’s modulus in either direction. During the experiment, the participant was seated in an adjustable chair with their finger placed directly under the substrate. A blindfold was used throughout the experiments to eliminate any visual cues. Within a pair, each stimulus was sequentially delivered to the participant’s index finger in a randomized order for a total of 20 trials. We also delivered the same stimulus twice to the finger to test the individual biases. After indented by two sequential stimuli, the participant was asked to report which of the two stimuli was more compliant, either the first or second. The switching time between two stimuli was about 2 s. In total, there were 4,000 indentations, consisting of 2 stimuli within a pair, 4 comparison pairs, 20 trials, and 25 participants for Experiments 1-4, with an additional 15 participants in Experiment 5. The average duration of this experiment per participant was about 80 min including breaks. We conducted the psychophysical experiments before biomechanical experiments for two reasons: one is to maintain a consistent time interval between two stimuli, as in biomechanical experiments the stimulus surface needs to be cleaned after each indentation (3-5 s); and the other reason is to ensure a greater cognitive attention from the participants.

#### Biomechanical experiments

During the experiment, the seven compliant stimuli noted above were delivered to the finger individually at a rate of 1.75 mm/s at 2 mm displacement, repeated 3 times per stimulus. Before indentation, each stimulus was brought into the finger with a light contact force (< 0.1 N) then slowly retracted to 0 N to ensure a consistency in initial contact states between trials. Any residual ink was removed from the stimulus surface after each indentation to attain the highest quality of imaging data. In total, there were 525 indentations, consisting of 7 compliant stimuli, 3 repetitions and 25 participants in Experiments 1-4. The average time to complete this experiment was about 70 min including breaks.

#### Skin stiffness modulation experiments

A new group of 15 participants were recruited for Experiment 5. To modulate their skin stiffness, a few drops of hyaluronic acid serum (Cosmedic Skincare Inc. Rocklin, CA), an agent that helps relax and hydrate the skin, was applied on the finger skin and allotted about 5 mins to dry at room temperature. We measured the participants’ skin stiffness and evaluated their perceptual performance before and after treatment using the approaches described above.

### Statistical analysis

To split the participants into subgroups based on their finger size, skin stiffness, and ridge breadth, we used a k-means clustering method. As shown for Experiments 1-4, clusters of size two were optimal using the elbow method in which the sum of squared distance between each point and the centroid in a cluster was plotted with number of clusters, Appendix A3. Principal component analysis (PCA) was then used to assess differences in the utility of skin deformation cues between the two groups. The total explained variance values from the first and second principal components for each cue were summed and compared for each group, Fig. 5(c).

Next, we employed multivariant regression model for each group to predict discrimination performance across participants based on their finger properties. The data were split into training (75%) and test (25%) sets. The independent variables were the change rate of five skin deformation cues and the dependent variable was discrimination rate from the psychophysical experiments. To validate the use of these regression models, we verified that the independent variables were not statistically dependent and the residuals were randomly distributed, Appendix A4. We used 10-fold cross validation to build subgroup models (n = 1,050 in the Group 1 model, n = 945 in the Group 2 model), and values of R^2^ and RMSE were used in evaluating model performance.

## ACKNOWLEDGMENTS

This work is supported in part by grants from the National Science Foundation (IIS-1908115) and National Institutes of Health (NINDS R01NS105241). The funders had no role in study design, data collection and analysis, decision to publish, or preparation of the manuscript.

## APPENDIX / SUPPLEMENTAL INFORMATION

**Appendix A1.**
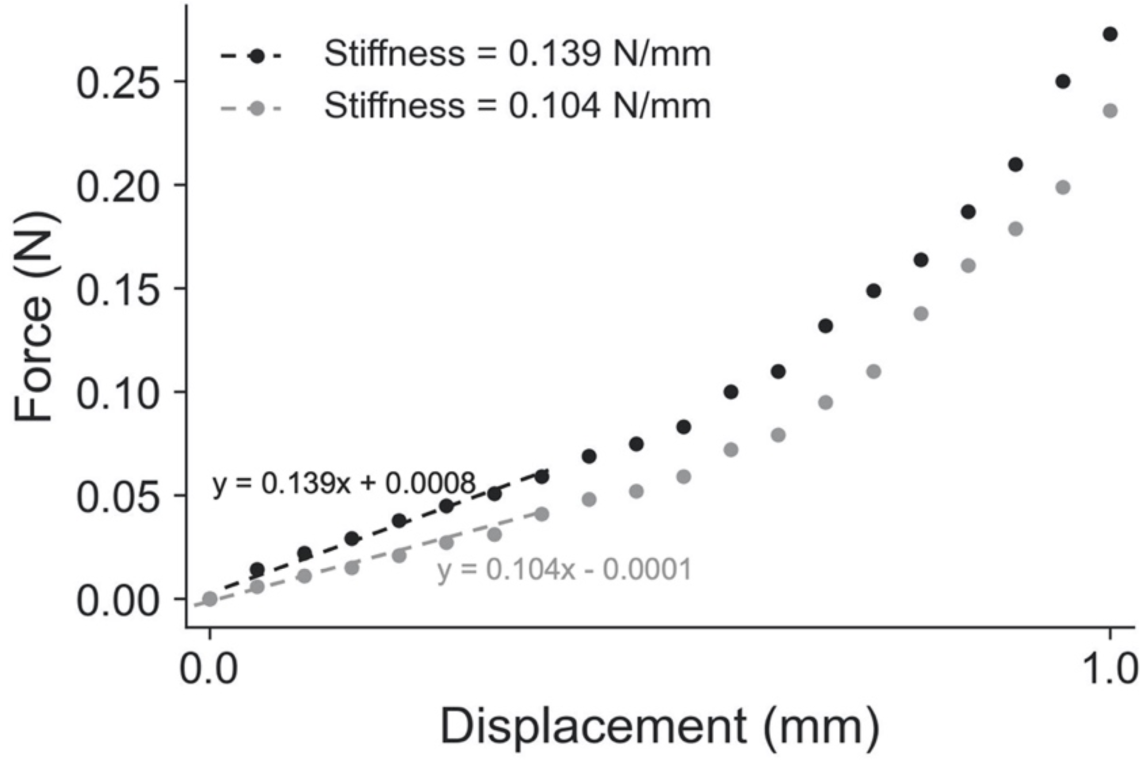
Example measurement of the finger skin stiffness of two participants. The light grey line (below) is participant 3, from Fig. 5(a), with softer skin, and the dark grey line (above) is participant 4 with stiffer skin. These example participants were selected due their similar finger size (237.9 mm^2^ for participant 3 and 234.4 mm^2^ for participant 4) but distinction in finger skin stiffness.

**Appendix A2.**
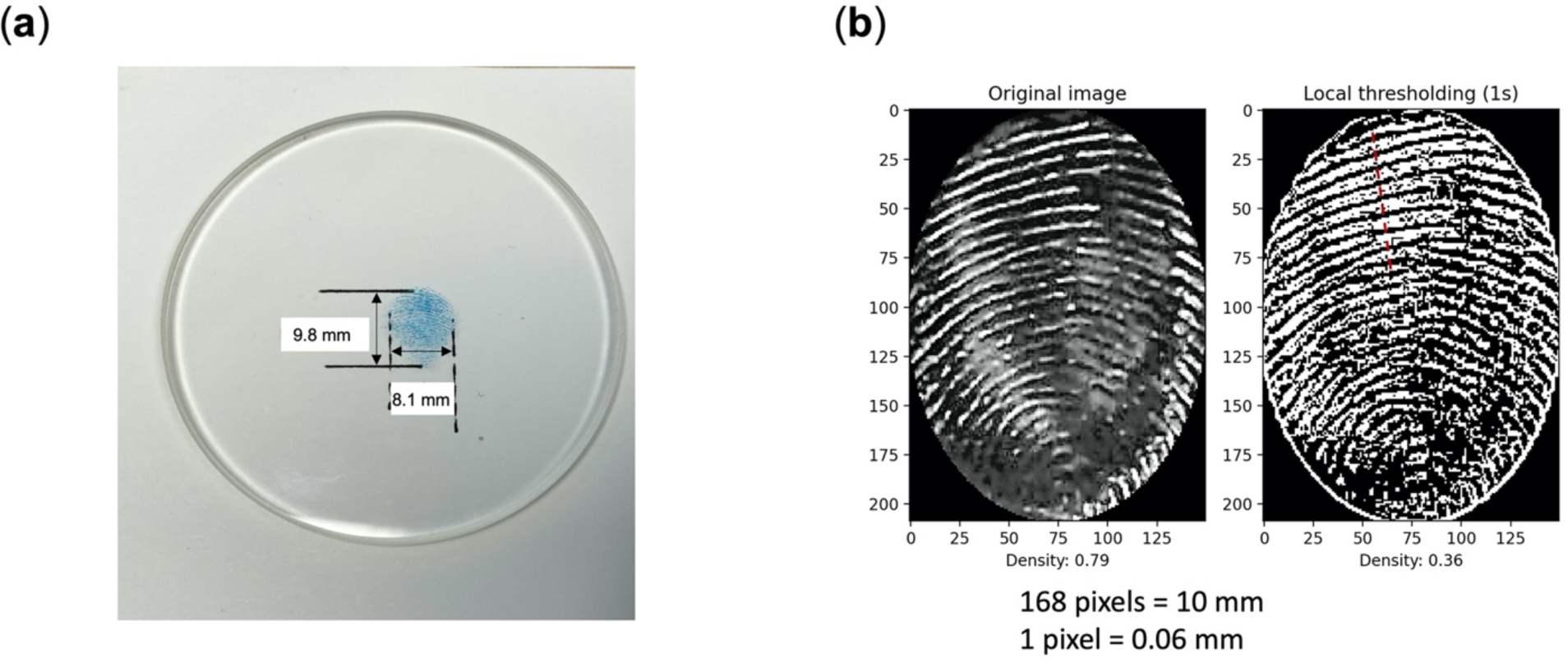
**Steps in measuring finger ridge breadth**. **(a)** Ink deposited contact region, deposited purposefully, between the finger skin and a glass plate from the compression test, with its length and width are measured by a digital caliper (resolution: 0.1 mm, accuracy: ±0.2 mm), which are the measurements used to obtain finger size. **(b)** Digital images of the same finger skin for use in obtaining fingerprint ridge breadth.

**Appendix A3.**
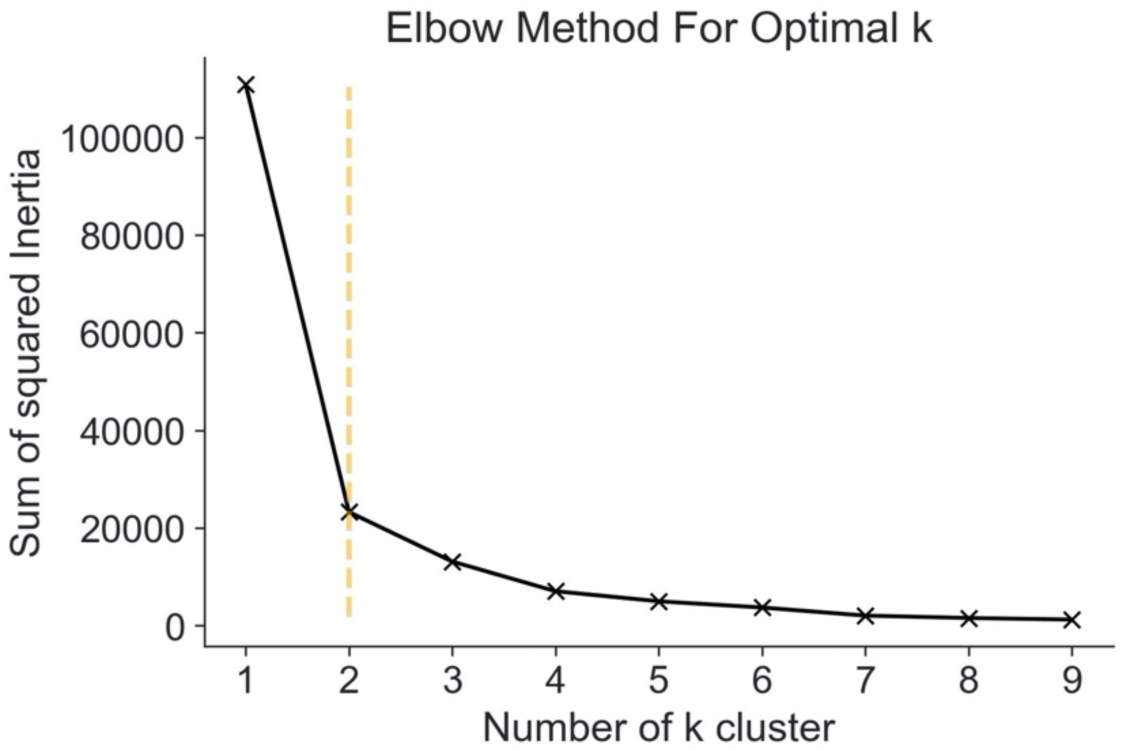
**Optimal number of clusters of study participants in K-means clustering method is k = 2**. The elbow method is used for determining the optimal number of clusters by calculating the sum of squared errors within clusters for different values of k and selecting the number for which change in the error first starts to diminish.

**Appendix A4.**
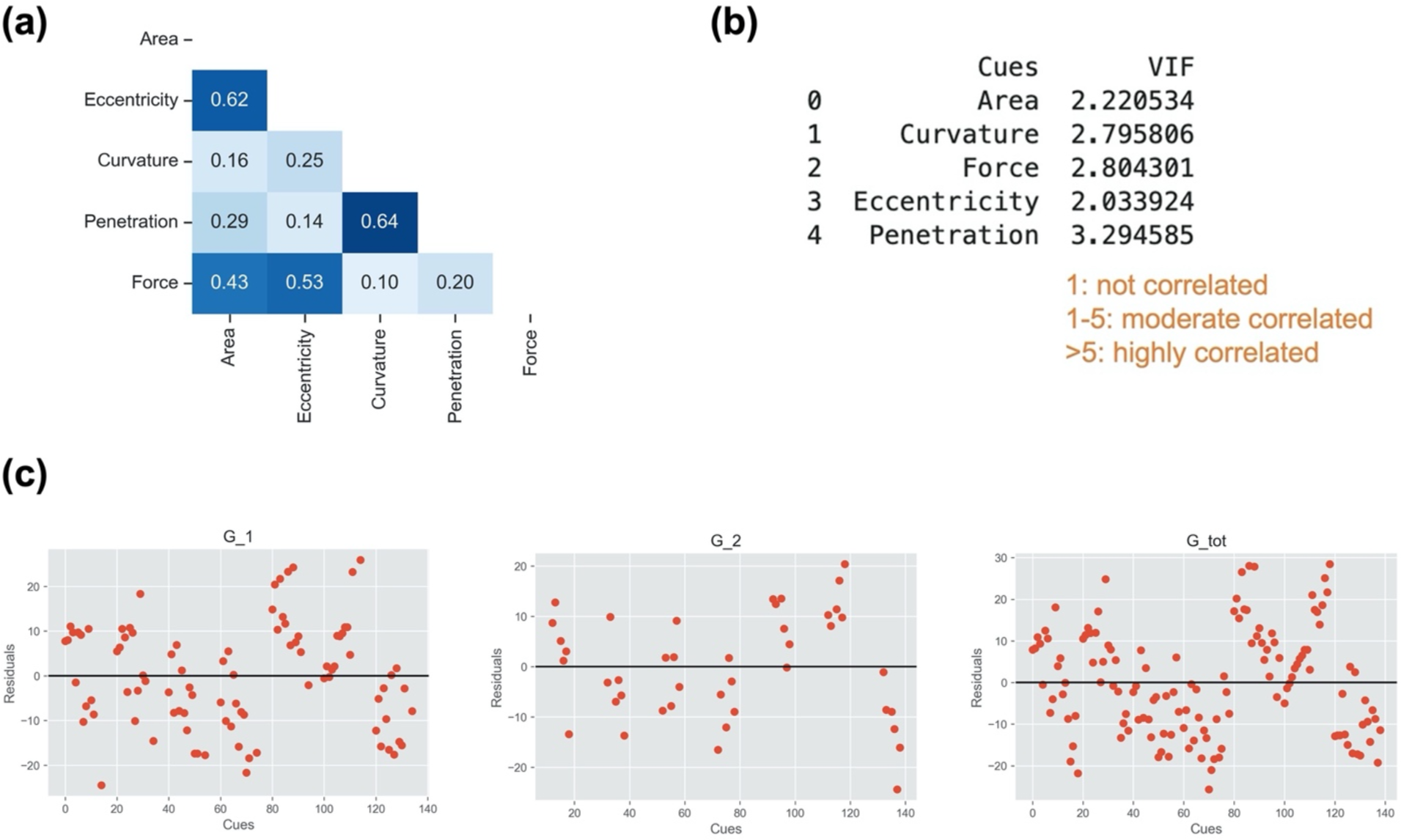
Validation of assumptions of correlation, covariance and distribution for the linear regression model. (a) For the 25 participants in Experiments 1-4 are shown the correlations between the independent variables, which describes how the relate with one another. The variables being the change rate of contact area, eccentricity, curvature, penetration depth, and force. In general, each variable provides unique value. **(b)** The covariance between the independent variables, which describes how each cue differs from each other. The numbers 1-5 are rules of thumb where if VIF is greater than 5, one should not use the regression model. **(c)** The residuals from the linear model are randomly distributed, which validates the use of the model.

**Appendix A5.**
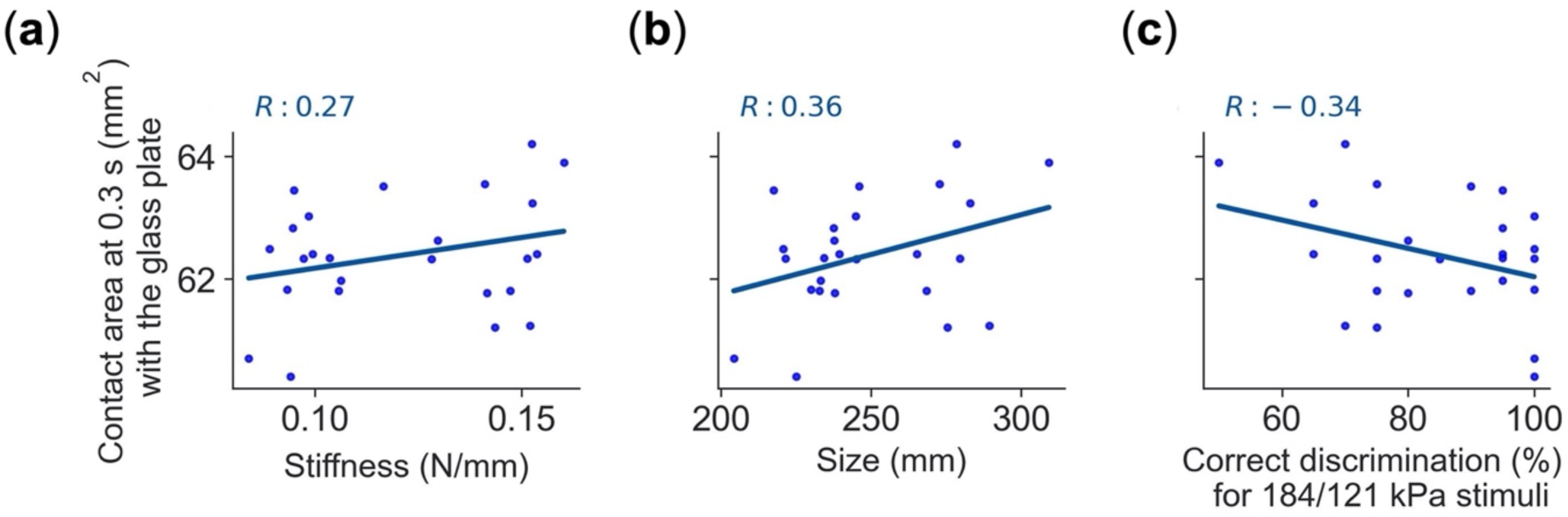
Larger fingers do not necessarily generate larger contact area, nor does greater contact area correlate with perceptual discrimination, or influence measurements of skin stiffness. Relationships across participants between contact area measurements at 0.3 sec indentation into the glass plate and finger stiffness, size, and with respect to correct discrimination for the least compliant pair of stimuli. **(a)** In the linear region of the force-displacement curve and right before stimuli become discriminable at 0.4 sec, i.e., at 0.3 sec indentation, skin stiffness has no significant effect on measurements of contact area. This indicates that the initial contact area does not significantly impact measurements of skin stiffness**. (b)** Finger size is also not correlated with initial contact area. This indicates that even for those participants with larger fingers, their contact area differs, which is likely due to individualized finger geometry. **(c)** Contact area measurements per participants are compared with his or her rate of correct discrimination for the least compliant pair of stimuli, again indicating a lack of correlation between the absolute value of contact area and perceptual ability.

